# Molecular Dynamics Analysis of Self and Microbial Peptides Bound to HLA-B Protein: A Multi-Parameter Framework

**DOI:** 10.64898/2026.02.14.705892

**Authors:** Sanju Singh

## Abstract

Molecular mimicry between pathogen-derived and self-peptides shown by MHC molecules is one of the critical mechanisms in the pathophysiology of autoimmune diseases. Numerous studied has been conducted in this field to identify sequence similarity, but evaluating structural and dynamic similarity, systematic computational frameworks remain limited. Therefore, we created an automated multi-parameter molecular dynamics analysis workflow and used it to compare three peptides (KP1, KP2, and KP3) generated from Klebsiella pneumoniae bound to HLA-B class protein with one human self-peptide (Annexin-derived, ANX). We assessed six complementing parameters using one microsecond-scale MD simulation: radius of gyration (Rg), solvent-accessible surface area (SASA), hydrogen bonding dynamics, MM-GBSA binding free energy, root mean square fluctuation (RMSF), and root mean square deviation (RMSD) to understand time-dependent structural and dynamic behaviour of all the peptide-HLA-B complex. Additionally, hydrogen bond occupancy and molecular mechanics generalised Born surface area (MM-GBSA) binding free energy calculations were performed to provide a more comprehensive assessment of complex stability. Our analysis suggests that KP1 exhibits structural features consistent with molecular mimicry, maintaining conformational stability, surface exposure, and interaction patterns comparable to ANX. In contrast, KP2 showed reduced stability, characterised by higher RMSD values and substantial hydrogen bond loss, whereas KP3 displayed intermediate behaviour, with relatively favourable energetics but noticeable conformational variability. Overall, the multi-parameter framework enabled differentiation among the candidate peptides based on combined structural, dynamic, and energetic properties. The workflow can be adapted for the analysis of larger peptide datasets and may provide a systematic approach for investigating potential autoimmune-relevant molecular mimics in microbial proteomes, with required adjustments according to the system.

## 1. Introduction

Autoimmune disorders have been identified as more than 100 chronic ^1^, complicated, and growing conditions in which the immune system has lost the capacity to discriminate between its own and alien components, due to which, immune responses are inappropriately directed against self-antigens, leading to tissue damage and the development of autoimmunity ^2^. This condition can happen due to a mix of genetic susceptibility and environmental triggers ^3^, and the common examples include rheumatoid arthritis (joints), type 1 diabetes (pancreas), multiple sclerosis (nervous system), systemic lupus erythematosus (multiple organs), psoriasis (skin), and ankylosing spondylitis ^4^. There is no single autoimmune gene, but certain genes increase risk, especially genes from the HLA (Human Leukocyte Antigen) family ^5^, which belong to the major histocompatibility complex (MHC) and help present antigens to immune cells ^6^. Some examples, such as the HLA-B27 gene is strongly associated with ankylosing spondylitis ^7^ and other immune-regulating genes, such as PTPN22, CTLA4, and IL23R, can also contribute to the same or other autoimmune conditions ^8^ ^9^.

Strong correlations have been observed by many investigations for HLA alleles, particularly the HLA-B family ^10^. These HLA-B class genes have been widely associated with inflammatory and autoimmune disorders like ankylosing spondylitis and associated spondyloarthropathiesand^11^. These correlations imply that T-cell recognition and antigen presentation play a key role in the development and progression of disease. ^12^. According to some studies, this involves the presentation of intracellular peptides to cytotoxic T lymphocytes by major histocompatibility complex (MHC) molecules, which shapes immune surveillance and self-nonself discrimination ^13^. Despite decades of investigation, the specific molecular processes by which the HLA-B class contributes to disease susceptibility are still not fully understood.

One proposed explanation for autoimmune disease is molecular mimicry, in which microbial peptides resemble host-derived self-peptides closely enough to trigger cross-reactive immune responses ^14^. In this mechanism, peptides derived from microbes, including those from the gut microbiota, may be presented by HLA molecules like endogenous antigens ^15^. This similarity can lead to unintended T-cell activation against self-proteins, and such cross-recognition may result in immune responses that target structurally related host proteins in addition to the pathogen, contributing to chronic inflammation and tissue damage ^16^ ^17^.

Numerous studies have been conducted that link microbial exposure to increased autoimmune susceptibility in genetically predisposed individuals, which extensively reported sequence similarities between HLA-B-restricted self-peptides and peptides derived from microbes present in our body. Many investigators connects some gut-microbiota such Klebsiella pneumoniae as a suspect, who derive mimicry peptides which influence cross reactivity ^18^ ^19^ ^15^. However, immune recognition is not governed by sequence similarity alone ^20^, it may also be influenced by factors such as protein–peptide dynamics, structural conformation, peptide–HLA binding stability and the surrounding immunological environment ^21^. Peptides with limited sequence similarity may adopt similar conformations when bound to HLA molecules, whereas peptides with higher sequence identity can exhibit distinct dynamic behaviour ^22^. Therefore, understanding autoimmune triggering requires integrating structural, dynamic, and functional analyses beyond simple sequence comparison, which relies primarily on static structural models, sequence alignment, or motif-based approaches ^23^.

Therefore, we address this gap in this study through comparative molecular dynamics (MD) analysis of one human self-peptide (Annexin-derived, ANX) and three *Klebsiella pneumoniae*-derived peptides (KP1, KP2, KP3) bound to HLA-B class. The objective of this work is to evaluate whether any of the microbial peptides exhibit structural and dynamic similarity to the self-peptide that could support a hypothesis of potential molecular mimicry. To this end, we systematically analyse multiple MD-derived descriptors, including root mean square deviation (RMSD), radius of gyration (Rg), root mean square fluctuation (RMSF), solvent-accessible surface area (SASA), number of Hydrogen bonds (H-Bonds) and relative Binding Energy (BE). These parameters collectively provide insight into peptide stability, flexibility, compactness, and interaction behaviour within the HLA binding environment.

By integrating these complementary analyses, we establish a multi-parameter computational framework for systematic comparison of self and microbial peptides in HLA-associated autoimmunity. The findings aim to identify microbial peptide candidates exhibiting ANX-like conformational characteristics and to provide a mechanistic basis for future structural, energetic, and immunological investigations into molecular mimicry-driven immune cross-reactivity.

## 2. Materials and Methods

### 2.1 Software and Computational Tools

All computational analyses were performed using open-source or freely available academic software. Bioinformatics preprocessing and automation were conducted using Python (version ≥3.10) with Biopython ^24^ for sequence handling and alignment. HLA-B class binding prediction was performed using NetMHCpan ^25^ ^26^ ^27^. Structural modelling and docking were carried out using AlphaFold-Multimer ^28^, while molecular visualisation was performed using PyMOL ^29^ and VMD ^30^. Molecular dynamics simulations were conducted using AmberTools23 ^31^ and GROMACS 2024.0 ^32^ ^33^ ^32^, and the Amber parameterised files were converted to GROMACS format using ParmEd 4.1.0 ^34^.

### 2.2 Computational Workflow Overview

This study employed an integrated computational pipeline to identify and characterise microbial peptides capable of molecular mimicry with a human self-peptide in the context of the HLA-B protein. The workflow comprised five major stages: (i) data preparation and peptide generation, (ii) HLA-B class binding prediction and sequence similarity screening, (iii) peptide-HLA molecular docking, (iv) molecular dynamics (MD) simulations, and (v) post-simulation trajectory correction and structural analysis. An overview of the workflow is provided in Figure 1.

**Figure 1.**
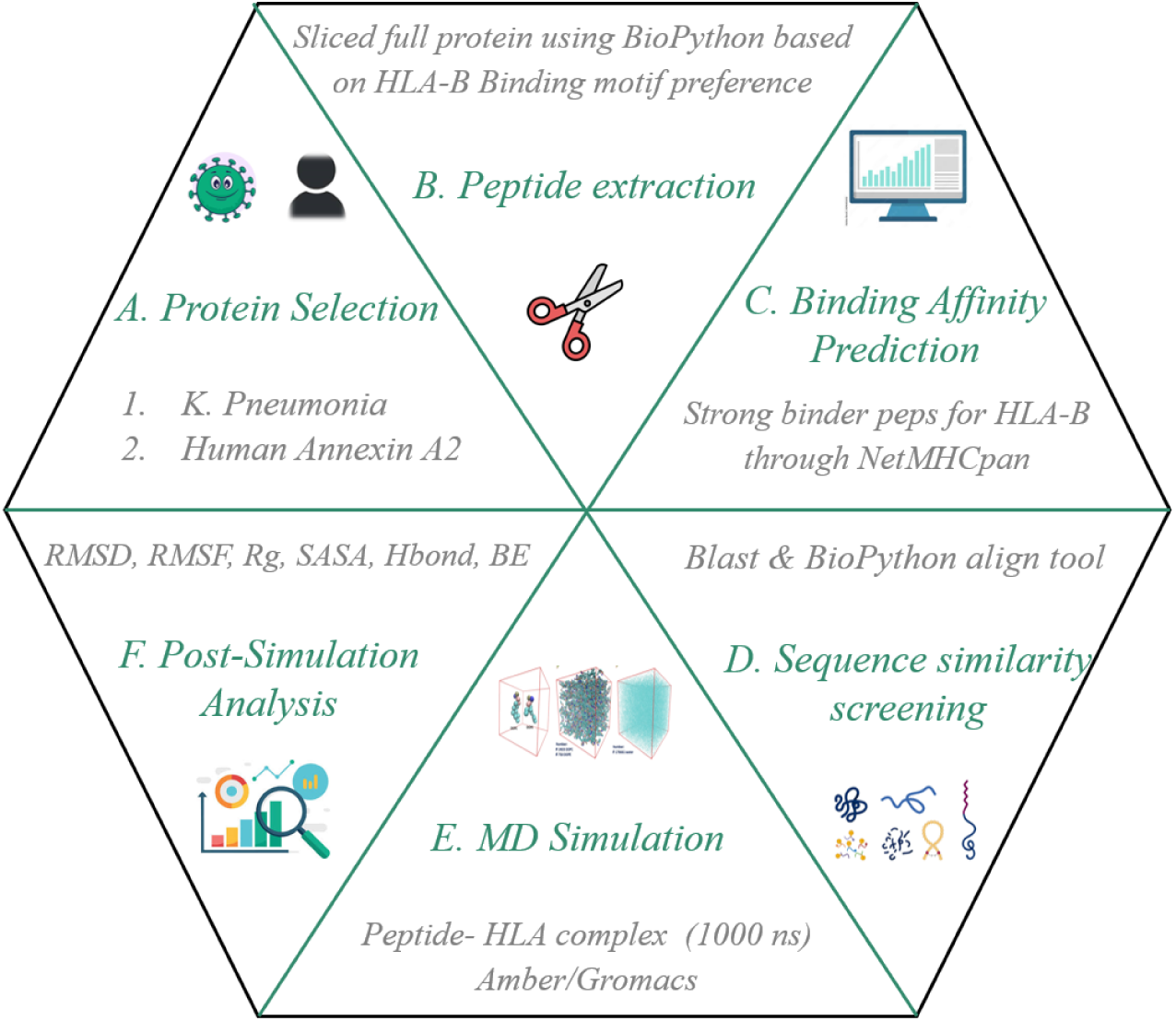
Schematic representation of the computational pipeline, including pairwise sequence alignment using Biopython, peptide modelling, and Molecular Dynamics (MD) simulation protocols.

### 2.3 Data Preparation

#### 2.3.1 Selection of Protein and Peptide Sources

To investigate peptide-level molecular mimicry relevant to HLA-B-associated autoimmunity, the HLA-B class protein crystal structure is obtained from the PDB data source (PDB ID: 5txs). On the other hand, two biologically relevant protein sources were selected for peptide generation: a human self-protein and a microbial proteome. Annexin (ANX) was selected as the representative human self-protein because of its well-documented roles in immune regulation and inflammatory signalling. Annexin influences bone cell activity, inflammation, and mechanotransduction processes central to abnormal bone formation and chronic inflammation in autoimmune diseases such as ankylosing spondylitis ^35^ ^36^. The full-length FASTA sequence of Annexin was retrieved from the UniProt (UniProt: P07355) database. As the microbial source, the complete proteome of *Klebsiella pneumoniae* was selected (Uniport: UP000000265) based on its widely reported association with HLA-B class-linked autoimmune disease and proposed role in molecular mimicry-mediated immune cross-reactivity ^37^.

#### 2.3.2 Peptide Generation & Peptide Slicing Strategy

Peptides of length 9-12 amino acids were generated from both human and microbial protein sequences, consistent with the known binding preferences of HLA class I molecules ^38^. Two complementary peptide slicing strategies were applied to ensure broad coverage while enriching for biologically plausible binders.

In the rule-free approach, peptides were generated using a sliding window without positional constraints, allowing unbiased exploration of the peptide space. In parallel, an anchor-rule-based strategy was employed to enrich for peptides containing known HLA-B anchor residue preferences, including positively charged residues at position 2 and hydrophobic or aromatic residues at the C-terminal position. ^39^ ^40^. Both strategies were applied consistently using custom Python script developed for this study to human and microbial sequences.

#### 2.3.3 Peptide Filtering and Similarity Screening

All generated peptides were screened for predicted binding affinity to HLA-B using NetMHCpan ^27^ and were classified based on predicted binding strength, and only strong binders were retained for downstream analysis. Approximately ten Annexin-derived peptides and ninety-two microbial peptides passed this filtering step. Later Sequence similarity between human and microbial peptides was evaluated using a two-stage approach. An initial similarity assessment was performed using BLAST-based comparison ^41^. Subsequently, quantitative pairwise sequence alignment was conducted using the Bio.Align the module of Biopython with the BLOSUM62 substitution matrix ^42^. Alignment scores were used to identify microbial peptides sharing conserved residues or motifs with human Annexin peptides.

Based on combined binding prediction and similarity screening finally total of sixty peptides (Ten human self-peptides and fifty microbial peptides) were finally identified. For each self-peptide, the top five microbial candidates were shortlisted to identify as potential mimicry partners. After this stage, three self-peptides and twelve microbial peptides (four microbial peptides for each one self-peptide) were taken to the next stage of molecular docking.

Peptide-HLA binding and presentation were predicted using NetMHCpan v4.1b (DTU Health Tech). Eluted ligand (EL) predictions were performed for the HLA-B27:05 allele, which is directly represented in the training dataset (distance = 0.000). Peptides with percentile rank ≤0.5 were classified as strong binders, while those with ranks between 0.5 and 2.0 were considered weak binders.

### 2.4 Molecular Docking

All the selected Peptide-HLA-B complexes were modelled using AlphaFold-Multimer via their respective web servers ^43^ as an initial structural screening step to obtain plausible peptide-HLA binding conformations. Each shortlisted 15 shortlisted Peptide–HLA complex models was generated using AlphaFold-Multimer, and the model with the highest-ranking confidence score was selected using a custom Python script. Based on docking quality and visual inspection, one human peptide (IRSEFKRKY) and three microbial peptides (GRSDFKGDY, GRSDFKGDYF, and SRSDRVAKY) were selected (Table 1) for molecular dynamics simulations as an initial pilot dataset to validate the computational workflow, establish quality control criteria, and generate preliminary insights. The validated pipeline will then be applied to the remaining peptide pairs to enable comprehensive mimicry screening in future work. Importantly, all the AlphaFold prediction files and associated confidence summaries used for model selection are available in the GitHub repository ^44^. Additionally, all the structural and stability interpretations and conclusions presented in this study are based exclusively on MD simulation analyses; the docking was just employed as an initial structural screening step to predict possible binding before long run MD.

**Table 1.** Peptide Sequences and Binding Predictions.

| Peptide | Source | Sequence | Length | %Rank | Affinity |
| --- | --- | --- | --- | --- | --- |
| ANX | Human ANXA2 | IRSEFKRKY | 9 | 0.12 | Strong |
| KP1 | *K. pneumoniae* | GRSDFKGDY | 9 | 0.19 | Strong |
| KP2 | *K. pneumoniae* | GRSDFKGDYF | 10 | 0.20 | Strong |
| KP3 | *K. pneumoniae* | SRSDRVAKY | 9 | 0.15 | Strong |

### 2.5 Molecular Dynamics Simulations

All molecular dynamics (MD) simulations were performed following a classical all-atom approach using AMBER and GROMACS software packages. Initial peptide-protein complex structures were prepared from docked models and processed using AmberTools. Protonation states at physiological pH (pH 7.0) were assigned using PropKa ^45^ and the web server PDB2PQR ^46^ ^47^, followed by structure cleanup using pdb4amber ^48^ to ensure compatibility with AMBER force fields.

System parametrisation was carried out using the AMBER ff14SB force field ^49^ for proteins and peptides. The complexes were solvated in an explicit TIP3P water box ^50^ with a minimum padding of 10 Å from the solute. Sodium and chloride ions were added to neutralise the system charge. All system preparation steps were performed using the tleap module of AmberTools. The AMBER-generated topology and coordinate files were converted to GROMACS-compatible formats using ParmEd. All subsequent simulations, trajectory processing, and analyses were conducted using the GROMACS software suite, executed on the Phase 2 DelftBlue supercomputer ^51^.

Energy minimisation was performed algorithm until a maximum force threshold of 1000 kJ mol⁻¹ nm⁻¹ was reached or for a maximum of 50,000 steps, using the steepest descent algorithm. The systems were then equilibrated under constant volume (NVT) conditions for 200 ps at 300 K using the velocity-rescaling (V-rescale) thermostat ^52^, followed by constant pressure (NPT) equilibration for 300 ps at 1 bar using the Parrinello-Rahman barostat ^53^. All the bonds were constrained using the LINCS algorithm, also involving hydrogen atoms ^54^, allowing a time step of 2 fs. Long-range electrostatic interactions were treated using the particle mesh Ewald (PME) method with a real-space cutoff of 1.0 nm and a Fourier grid spacing of 0.16 nm. Van der Waals interactions were truncated at 1.0 nm, and periodic boundary conditions were applied in all three dimensions ^55^. Production MD simulations were carried out for 1 μs under NPT conditions at 300 K and 1 bar. Coordinates were saved every 10 ps in compressed trajectory format for subsequent analyses.

### 2.6 Post-Simulation Trajectory Processing

Raw MD trajectories obtained after a complete MD run were corrected to remove periodic boundary conditions (PBC) artefacts using the whole and nojump options implemented by gmx trjconv, followed by centring of the protein-peptide complex initial structure within the simulation box. For visualisation and comparative analyses, trajectories were rotationally and translationally fitted to the protein backbone (Figure 3A). Additional reduced trajectories were generated by systematic frame skipping to facilitate long-timescale inspection ^56^.

### 2.7 Post-MD Analysis

After all the simulation process the stability and flexibility of the structure were assessed using RMSD and RMSF analyses, while the compactness and solvent exposure were evaluated using radius of gyration and solvent-accessible surface area (SASA) calculations. Peptide-HLA interaction persistence was examined through hydrogen bond analysis. Relative binding free energies (BE) were estimated using the Generalised Born and Surface Area continuum solvation method (MM/GBSA) ^57^ ^58^. Trajectory visualisation was performed using VMD and PyMOL.

### 2.8 Workflow Automation and Reproducibility

The computational pipeline was designed to build an automated, scalable and repeatable execution workflow which can use customised Python scripts, procedures and follow steps to conduct HLA binding prediction, similarity screening, MD simulation setup, and trajectory analysis. To process a large number of candidates efficiently, NetMHCpan-based binding prediction and peptide slicing from complete proteomes were parallelised. On the other hand, to ensure consistency across various peptide-HLA complexes and facilitate batch processing, we used the same scripted workflow, differing only in the peptide sequence and identifier and setup (solvation, neutralisation, energy minimisation, and equilibration). Post-simulation analysis scripts were used to extract common structural descriptors (RMSD, RMSF, SASA, radius of gyration, hydrogen bonds, and MM-GBSA estimates) and to generate required plots and summary tables.

All scripts, some parameter files, images and tables are made available in the project’s public GitHub repository ^44^ with accompanying documentation provided via Jupyter Book ^59^. While computational results may depend on software versions and hardware environments, the provided resources allow the workflow to be reproduced in principle and adapted by other researchers with slight modifications according to their work requirements. The pipeline was tested on the present dataset and may be readily extensible for the larger set of peptides and additional HLA alleles.

## 3. Results and Discussion

To explore potential structural and dynamical similarity between a host-derived self-peptide (ANX) and selected microbial peptides (KP1, KP2, KP3), atomistic molecular dynamics simulations were analysed using multiple complementary descriptors. The data reported after analysis are primarily intended to compare and rank the examined peptides under consistent computational settings. As a result, the given measures were utilised to demonstrate relative similarity in interfacial stability, persistence behaviour, and binding potential, while supporting overall performance levels. Structural characteristics such as RMSD, radius of gyration, and SASA were measured at 10 ps intervals throughout the 1 µs trajectories. The initial 500 ns were set aside for equilibration, and time-averaged values, standard deviations, and observed ranges were computed from the remaining production phase (500–1000 ns) for each peptide-HLA complex ^60^. These methods allowed us to exclude pre-equilibration values from the production phase and allowed a uniform comparison across all the systems for a systematic comparison of microbial peptides to the self-peptide reference.

### 3.1 Structural Stability and Equilibration Dynamics

#### Backbone RMSD Analysis

The RMSD analysis was determined to assess structural stability as well as maintenance of initial binding postures during simulation for each peptide-HLA complex ^61^. In general, higher RMSD values indicate more conformational variation, whereas stability within a given range indicates the achievement of dynamic equilibrium ^62^.

The RMSD of the ANX-HLA complex rose during the simulation and quickly stabilised within a small range, primarily below 0.5 nm. The mean value of RMSD was 0.41 ± 0.05 nm. The graph achieved a steady plateau following equilibration and without showing any noticeable peaks, which indicates that the initial binding conformation and overall structural integrity were maintained throughout the trajectory with a stable and lower RMSD value ^63^. A similar equilibration pattern was seen for the KP1-HLA complex that followed an initial adjustment phase and then stabilised after around 25 ns with minor fluctuation within a similar low range. The stable plateau in the plot indicates that KP1 has a stable binding mode with only small conformational changes, while the absence of a continuous ascending trend contributes to the trajectory’s convergence. The computed mean RMSD for KP1 is 0.47 ± 0.02 nm, which was very close to that of ANX and had a similar parallel trend in the plot, which can be assessed as similarity in the binding poses and conformational variations.

In contrast, the KP2-HLA complex showed a significant rise in RMSD, followed by stabilisation at a higher plateau (∼1.52-1.62 nm), with a mean RMSD of around 1.56 ± 0.02 nm. This large variance shows that extensive conformational rearrangement may occurred in complex structure before reaching equilibrium ^64^. Whereas, the higher plateau indicates that it may have physically different and more flexible binding conformation than ANX and KP1, which can be also seen on figure 2A. On the other hand, KP3-HLA complex demonstrated intermediate behaviour throughout the simulation, with an average RMSD value of approximately 0.74 ± 0.04 nm. Although a stabilising tendency was observed, minor fluctuations persisted throughout the trajectory up to 0.9 µs, suggesting that additional simulation time may further clarify the extent of convergence and plateau formation ^65^. While the system also does not show continuous and larger divergence, the only minor fluctuations indicate a slightly dynamic binding mode relative to the more tightly stabilised ANX and KP1 complexes.

**Figure 2:**
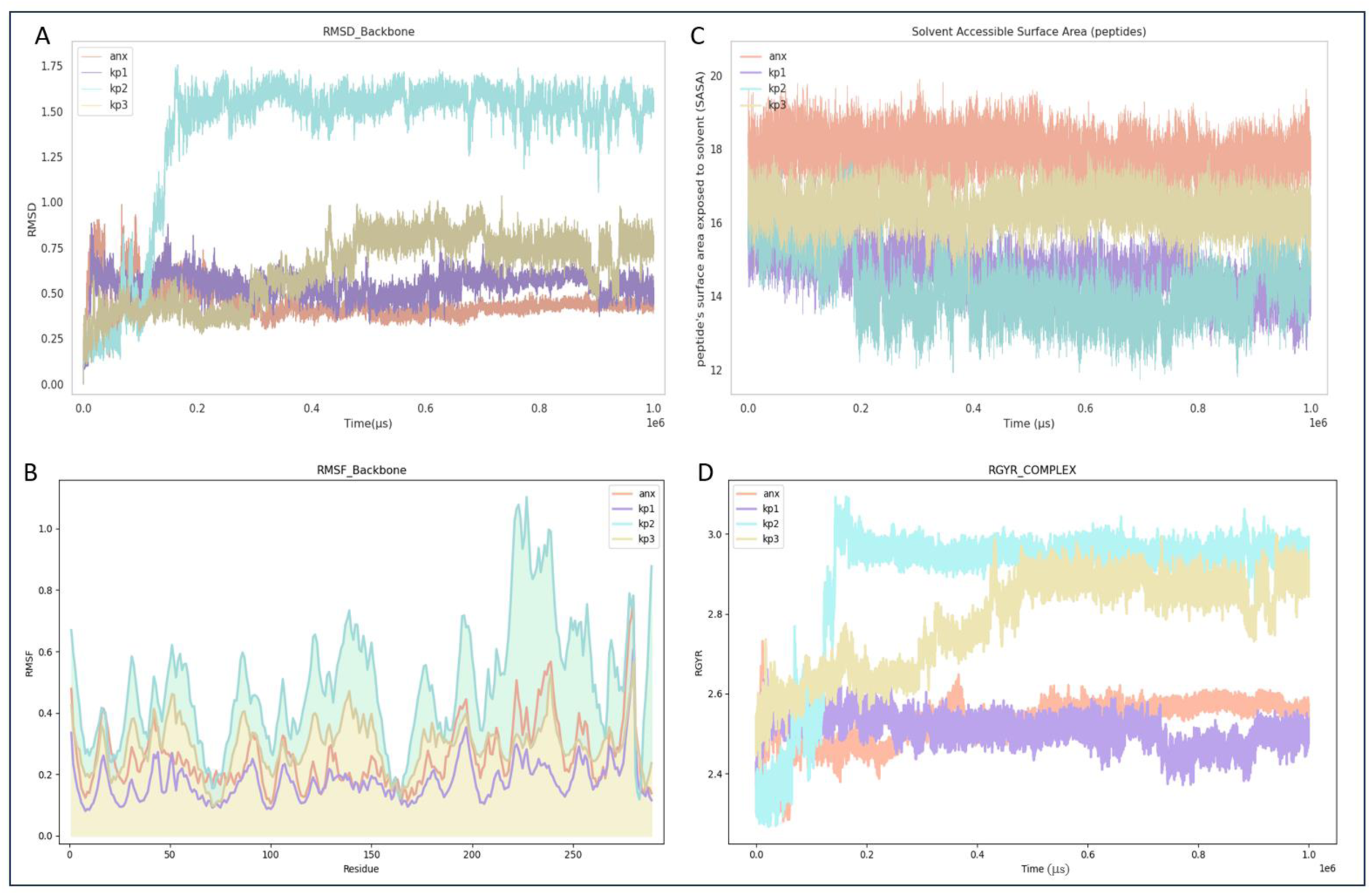
Structural stability and conformational changes of peptide complexes during molecular dynamics simulation: Analysis of key structural metrics across the 1 microsecond simulation trajectories for human Annexin (ANX) and microbial peptides (KP1, KP2, KP3). (A) Showing Root Mean Square Deviation (RMSD) Plot of backbone of complex. (B) Per residue flexibility Root Mean Square Fluctuation (RMSF) plot. (C) Solvent Accessible Surface Area (SASA): peptides surface area exposed to solvent. (D) Radius of Gyration (Rg) plot for complex. Colors of the plots used for individual peptides: Orange-ANX (anx), Purple-KP1(kp1), Cyan-KP2 (kp2), Yellow-KP3 (kp3).

Overall, comparative analysis of RMSD trends indicates that ANX and KP1 achieve rapid equilibration and sustained structural stability, and KP3 displays intermediate conformational variability; in contrast, KP2 undergoes substantial rearrangement before stabilisation at a higher deviation. These observations highlight differences in dynamic behaviour among the peptide-HLA complexes while emphasising highly comparative trajectory patterns between ANX and KP1 with similar equilibration trends and close RMSD value ^66^ ^67^.

#### Residue-Level Flexibility (RMSF)

The RMSF study determines the fluctuation of protein during the MD simulation to confirm the flexible regions of the protein that can reveal the local dynamic behaviour of the protein backbone. ^68^, and can provide major insights into how different peptides influence the flexibility of specific HLA regions ^69^. A higher RMSF value indicates that the atoms in the molecule have a greater change in position during the simulation, suggesting that these atoms are more flexible. On the contrary, a lower RMSF value indicates that the atoms have fewer positional changes, suggesting that these atoms are more stable during the simulation. ^70^ ^71^. Figure 2B represents the RMSF plot for all four complexes across the entire HLA protein sequence.

For the ANX-HLA complex, the backbone RMSF values remained mostly between 0.1 and 0.5 nm, with the average RMSF value around 0.30 ± 0.24 nm. The lower fluctuations were observed in the helix regions (around residues 55-85 and 135-175), indicating that the core binding groove remained relatively stable during the overall simulation. while slightly higher fluctuations can be seen at the N- and C-terminal ends, which is expected as terminal regions are generally more flexible than the others ^72^. Similarly, KP1-HLA complex showed a fluctuation pattern which closely resembles that of ANX with slightly lower RMSF values, approximately ranging from 0.08 to 0.4 nm. The helix regions (55-85 and 135-175) remained similarly stable, as shown in ANX, and similar higher fluctuations were observed near the terminal regions. However, the KP2-HLA complex displayed noticeably higher RMSF values, averaging around 0.77 ± 0.15 nm, with fluctuations increasing by approximately 0.4-0.5 nm compared to ANX and KP1 complexes. Although reduced fluctuations were noted in segments 65-75 and 160-165 (ranging approximately between 0.20-1.12 nm), the broader RMSF distribution indicates different dynamic behaviour relative to the other complex’s specifically with the reference one. This observation suggests that local stabilisation alone may not be sufficient to maintain a self-like flexibility profile necessary for potential cross-reactivity ^73^.

The KP3-HLA complex showed intermediate behaviour in which the RMSF values were higher than KP1 but lower than KP2 (0.09-0.56nm) with similar higher fluctuation near the terminal regions. The fluctuation pattern in the HLA backbone appeared somewhere closer to ANX but not identical with the average RMSF value 0.33± 0.24, which is almost very near to ANX’s RMSF value. Additionally, the core binding regions remain rather stable on a very limited region, with considerable decreases within residues 70-75 and 160-170, indicating stable tiny binding grooves in the complex structure.

Collectively, the data of the RMSF profile reveal that both ANX and KP1 have a similar and stable flexibility pattern, especially in the important helical regions that form the peptide-binding groove. In comparison, KP2 shows higher backbone movement, which suggests more flexible or altered structural behaviour within the complex. However, KP3 shows moderate behaviour; it is somewhat similar to ANX but does not have a completely identical plot. Overall, the findings indicate close resemble behaviour of the KP1 to that of the reference peptide, whereas KP2 have adopted a more flexible and structurally different conformation, while KP3 appears to behave somewhere between these two systems.

### 3.2 Solvent-Accessible Surface Properties and Global Compactness

#### Peptide SASA Dynamics

Solvent accessible surface area (SASA) was calculated for each peptide to evaluate the extent of solvent exposure and infer differences in binding compactness within the HLA groove, as it can predict the extent of exposed molecular surface of the molecule available for immune recognition ^74^ ^75^. Given Figure 2C represents the plots for fluctuation in SASA values during simulation, and the average SASA value of each complex is shown by Table 2. It can be seen that ANX and KP3 exhibit relatively higher average SASA values, centred around ∼16-18 nm², with closely stable profiles throughout the simulation whereas, KP1 and KP2 display comparatively lower SASA values, ranging approximately between ∼13-15 nm², accompanied by noticeable temporal fluctuations.

**Table 2.** Comparative Structural and Energetic Analysis of Peptide-HLA-B27 Complexes.

| Parameter | ANX (self) | KP1 | KP2 | KP3 |
| --- | --- | --- | --- | --- |
| RMSD (nm) | $0.41 \pm 0.05$ | $0.47 \pm 0.02$ | $1.56 \pm 0.02$ | $0.74 \pm 0.04$ |
| RMSF (nm) | $0.30 \pm 0.24$ | $0.22 \pm 0.15$ | $0.77 \pm 0.15$ | $0.33 \pm 0.13$ |
| SASA (nm<sup>2</sup>) | $17.71 \pm 0.002$ | $14.77 \pm 1.09$ | $14.93 \pm 0.51$ | $16.24 \pm 0.05$ |
| Rg (nm) | $2.51 \pm 0.05$ | $2.49 \pm 0.02$ | $2.98 \pm 0.00$ | $2.85 \pm 0.06$ |
| H-bonds (average) | $17.0 \pm 0.0$ | $14.8 \pm 2.1$ | $6.7 \pm 1.5$ | $18.0 \pm 2.3$ |
| $\Delta G$ (kcal/mol) | $-105.5 \pm 7.2$ | $-89.3 \pm 20.3$ | $-37.6 \pm 10.7$ | $-88.9 \pm 18$ |
| $\Delta\Delta G$ vs ANX (kcal/mol) | 0.0 | 16.2 | 67.9 | 16.6 |
| Mimicry Assessment | Reference | Strong | Weak | Intermediate |

RMSD: backbone root mean square deviation; RMSF: root mean square fluctuation; SASA: solvent-accessible surface area; Rg: radius of gyration; $\Delta G$ : MM-GBSA binding free energy; $\Delta\Delta G$ : relative binding free energy. Mimicry assessment: Strong ( $\geq 5/6$ parameters concordant), Intermediate (3-4/6), Weak ( $< 3/6$ ).

Primarily, with a mean peptide SASA of roughly 17.71 ± 0.002 nm^2^ and stayed comparatively constant throughout the trajectory, which suggests that there was no significant dissociation or unfolding and it showed only moderate solvent exposure ^76^. In contrast, KP1 displayed a lower mean peptide SASA of roughly 14.77 ± 1.09 nm^2^, suggesting a relatively lower solvent exposure, which is in line with the its stable RMSD and RMSF profiles similar to reference peptide. This finding implies that KP1 can adopt a compact binding conformation inside the groove and accommodated more tightly without showing signs of structural distortion as ^77^ shown in Figure 2A.

Remarkably, the KP2-HLA complex exhibited a comparatively lower peptide SASA (average around 14.93 ± 0.51 nm²) despite elevated average RMSD and RMSF values (Table 2). As we can see in the SASA profile, the peptide’s exposure was slightly higher at initial state of simulation, but it significantly decreased after about 0.2 μs. This reduction may correspond to a structural rearrangement observed during simulation, where the peptide partially entered the groove and adopted a half-bound conformation (self-association), then formed a loop-like structure while remaining partially exposed from the pocket, as shown in Figure 3A. This could be the reason of remark ably low SASA value compared to the other peptides ^78^ ^79^. Although there was a slight increase in SASA towards the trajectory’s later phase, the values remained primarily on the lower side. This indicates the main cause of fluctuating but predominantly decreased value of SASA can be due to conformational rearrangement and compaction, rather than full groove accommodation or improved structural stability ^78^.

**Figure 3:**
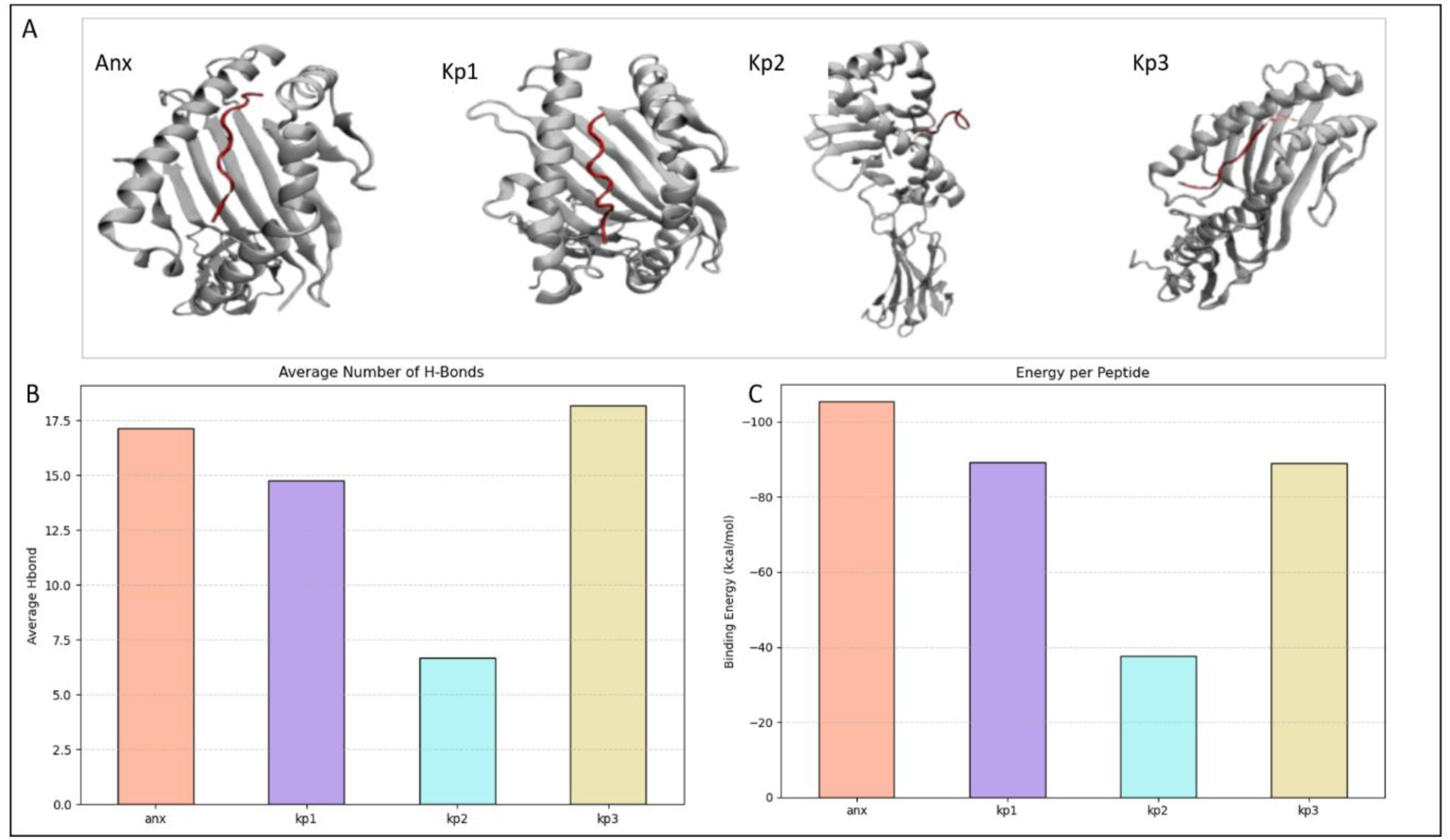
Intermolecular interactions and binding energy analysis of protein-peptide complexes: Visual representation and quantitative analysis of the interactions between the Annexin protein (Anx) and microbial peptides (kp1, kp2, and kp3). (A) Representative molecular structures of the four complexes shown in their final binding modes after molecular dynamics simulations. The protein is shown in a grey ribbon representation, with interacting residues highlighted in red. (B) Average number of intermolecular hydrogen bonds formed between the Annexin protein and the respective peptides throughout the simulation trajectory. (C) Calculated average binding free energies (kcal/mol) for each complex.

The mean peptide SASA of the KP3, on the other hand, was closer to that of the ANX, at about 16.24 ± 0.05 nm², which implies that the binding conformation is retained but moderately exposed towards the solvent. We observed some later fluctuation in the RMSD plot of KP3, but the SASA profile shows that the peptide does not collapse or dissociate extensively from the system, instead remaining intact during the simulation (Figure 3A).

Overall, it can be assessed that the SASA analysis itself is not a direct indicator of structural stability ^80^, as we can see the SASA values of KP1 and KP2 are similarly lower, but their dynamic behaviours are very different while more dynamically accommodated and naturally exposed binding mode within the groove may be the reason for the relatively higher SASA values shown by ANX and KP3. These findings highlight that relying on SASA magnitude alone to assess binding and stabilization quality for mimicry potential is not sufficient, it’s important to integrating SASA with RMSD and RMSF analyses to distinguish between real structural stabilised compact conformation and some varied conformation not related with the actual objective of work ^81^.

#### Complex Radius of Gyration (R_g_)

The radius of gyration (Rg) was calculated to examine the compactness and overall structural stability of the HLA-peptide complexes during the simulation. ^82^. Rg reflects the folding state of the protein-peptide system, where a relatively constant value indicates a stable and compact structure, while continuous variation may suggest structural expansion or rearrangement. ^70^ ^83^. The summarised average Rg values and the comparative graphical representation of all complexes are shown in Table 2 and Figure 2D, respectively. The reference peptide ANX showed an average Rg value of around 2.53 ± 0.03 nm (Table 2), indicating a stable and reasonably compact conformation throughout the simulation with minor fluctuations, which are observed, mainly within the range of 2.50-2.55 nm, suggesting its consistent structural compactness and folding ^83^.

Similarly, KP1 exhibited mean Rg value of 2.49 ± 0.02 nm (slightly lower than ANX), indicating marginally greater compactness compared to ANX. The complex fluctuated within a narrow range of 2.45-2.55 nm and stabilised early; its pattern on the plot closely resembled that of ANX for most of the simulation period, suggesting a similar level of structural compactness between both the complexes ^60^. In contrast, the higher plot of KP2 indicates reduced compactness compared to ANX and KP1, though it achieved equilibrium early in the simulation and displayed limited fluctuations after stabilisation, its overall Rg remained consistently higher, at 2.95 ± 0.03 nm. These observations suggest that the KP2-HLA complex adopts a more expanded conformation, which is consistent with its higher RMSD and RMSF values (Table 2), which indicates greater conformational flexibility in the complex. Similarly, KP3 exhibited an average Rg value of 2.87 ± 0.04 nm, which is also higher than ANX and KP1 fluctuated between 2.70 and 2.90 nm, indicating moderate compactness but comparatively lower than the reference system ^75^.

Overall, the Rg analysis suggests that ANX and KP1 maintain comparable compactness and structural stability, whereas KP2 and KP3 adopt relatively fewer compact conformations. The close similarity between ANX and KP1 in both average Rg values and fluctuation patterns supports the observation that KP1 maintains a structural behaviour more similar to the reference peptide within the HLA binding groove.

### 3.3 Intermolecular Interaction Networks Hydrogen Bond Analysis

Hydrogen bonding between peptide and HLA-B residues is a key factor that decides the specificity and stability of the binding present in the docked complex ^84^ ^85^. The average number of hydrogen bonds that formed between each peptide-HLA complex during the simulation is given in Table 2, and the comparative graph showing H-bond numbers is shown in Figure 3B. It can be seen that a strong interaction network that stabilises peptide binding was formed by the self-peptide ANX, which formed an average of 17 H-bonds throughout the simulation.

On the other hand, KP1formed around 14.8 ± 2.1 H-bonds, which accounted for 86% of the self-peptide interaction density and showed a significant number of interactions for higher stability of the peptide inside the binding groove. Additionally, this high H-bonding capacity retention can imply to the condition that KP1 might interact with the same residues as ANX, which can also be noticed by similar stability regions in RMSF plots as well as via per-residue binding energy values ^84^, as shown in Table S2. Notably, KP3 exhibited the highest average H-bond count, nearly 18.0 ± 2.3, which is unexpected due to its elevated Rg fluctuations. This unexpected result might suggest compensatory interactions that stabilise a binding geometry that would otherwise be less optimal ^86^. Interestingly, KP2 formed substantially fewer hydrogen bonds (6.7 ± 1.5), corresponding to less than 50% of the interaction network observed for all other peptides. This profound deficit in hydrogen bonding correlates directly with the elevated RMSD, increased R_g_, and reduced SASA ^87^ due to loop formation in the peptide observed in Figure 3A for KP2.

### 3.4 Binding Thermodynamics and Energetic Mimicry

#### MM-GBSA Binding Free Energy

To assess the relative binding affinities of the microbial peptides, MM-GBSA calculations were performed on equilibrated trajectory snapshots. ^88^ ^89^. While MM-GBSA is not suitable for absolute affinity prediction, it provides reliable relative rankings for comparing similar peptides binding to the same MHC allele^16^. As shown in Figure 3C, for implicit solvation models applied to peptide-protein complexes, absolute binding free energies were elevated compared to experimental measurements as expected ^90^ ^91^. However, Relative binding free energy differences (ΔΔG) provide meaningful rankings for peptide comparison (refs). The self-peptide ANX exhibited ΔG = −105.49 ± 7.2 kcal/mol. Microbial peptides KP1 (−89.30 ± 20.32 kcal/mol) and KP3 (−88.88 ± 18.00 kcal/mol) showed similar energetic profiles with moderate deviations from ANX (ΔΔG = 16.2 and 16.6 kcal/mol, respectively), indicating thermodynamic compatibility. In contrast, KP2 (−37.60 ± 10.7 kcal/mol) exhibited a 67.9 kcal/mol deficit, indicating substantially weaker binding.

The per-residue energy decomposition analysis, as shown in Table S2, demonstrates how the interactions between molecules work and helps to understand why some peptides are better at mimicking others ^88^ ^58^. The result revealed that the self-peptide ANX can bind mainly due to amino acids present at position 2 (Arg) and position 9 (Tyr), with contributing overall Binding energy −10.14 kcal/mol and −13.27 kcal/mol, respectively. ANX also significantly interacts with residues Glu64, Lys147 and Trp148 of the HLA-b molecule, which contribute −5.87 kcal/mol, −5.07 kcal/mol and −4.64 kcal/mol of energy, respectively. Similarly, the Peptide KP1 preserves this energetic signature residues with nearly identical contributions from position 2 (Arg: −10.54 kcal/mol) and position 9 (Tyr: −10.42 kcal/mol), which demonstartes a faithful reproduction of the anchor residue interaction network that defines the self-peptide binding mode. Along with that, the HLA-side interactions are similarly maintained in the KP1-HLA complex, with Glu64 (−3.04 kcal/mol) and Lys147 (−8.73 kcal/mol) providing consistent stabilisation and close functionality.

Additionally, peptide KP3 also exhibit similar interaction in both anchor positions 2 and 9, with estimated BE around −10.8 kcal/mol and −11.5 kcal/mol, respectively, very close to the reference peptide. It also shows comparable binding energy due to interaction with HLA_B residues Glu64 and Arg63 at around −3.22 and −3.02, respectively, which suggest comparable interaction to that of KP1 and ANX. Due to this observation, we can also suspect peptide KPKP33 as a mimicry candidate with a moderate possibility due to its slightly different RMSD and Rg plots. Noticeably, the KP2 has a very different C-terminal anchor interaction. At position 9 (Tyr), it is only contributing a −1.58 kcal/mol, that too not due to interaction with protein, as can be seen during the simulation (Figure 3A), although KP2 is contributing at position 2 (Arg: −11.09 kcal/mol), the C-terminal anchor is extremely important for stable binding, which is completely ablated here. In addition, the HLA-B residues Lys147 and Trp148 also contribute significantly lower than the other peptides (−0.62 and −0.57 kcal/mol, respectively).

On the other hand, KP2 is still doing okay with the position 2 anchor, which is Arg and has −11.09 kcal/mol. Because the C-terminal anchor is not working well, the whole peptide-binding register is not stable. We can see this because the HLA residues, like Lys147, are not contributing much as they should be. The energy for ANX is −0.62 compared to −5.07 kcal/mol. The energy for Trp148 is −0.57 compared to −4.64 kcal/mol.

This tells us about the energy of each part of KP2. The overall BE of KP2 ( ̴ −37.6 Kcal/mole))is very weak (ΔΔG 67.9 kcal/mol), which clearly suggests KP2 fails to engage the essential anchor residue network, is unable to form a stable and high-affinity binding complex and is incapable of supporting molecular mimicry.

### 3.5 Integrated Multi-Parameter Mimicry Assessment

As summarised in Table 2, KP1 demonstrates remarkable consistency with the self-peptide ANX across all parameters and maintains stable RMSD trajectories, also preserves native RMSF patterns, exhibits similar and compact R_g_ plots and retains substantial hydrogen bonding capacity (86% of ANX) as well as achieves comparable binding free energy (ΔΔG = 16.17 kcal/mol). On the other hand, Peptide KP3 presents an intermediate case with similar SASA value and even more hydrogen bonds, although it exhibits conformational heterogeneity. The observation is concluded due to variable R_g_, slightly elevated RMSD and RMSF, clearly KP3 may adopt self-like conformations but lacks the conformational stability to consistently maintain them.

However, KP2 fails across all structural and energetic criteria covered in the analysis with weakened binding (ΔΔG = 67.59 kcal/mol), reduced H-bonding (less than 40% of ANX), elevated structural deviations (RMSD, R_g_, RMSF), and dramatically reduced SASA. Although it is important to note that this computational analysis has some limitations, as it can only provide structural and dynamic evidence based on computational models for mimicry potential, it does not directly measure immunological cross-reactivity as measured in wet lab. Additionally, TCR recognition depends not only on peptide-MHC structure but also on TCR repertoire diversity, antigen processing efficiency, and immunoregulatory context ^92^ ^93^. Nevertheless, the strong multi-parameter concordance observed for KP1 can provide a computational foundation for prioritising this peptide for experimental validation

## 4. CONCLUSION

The present study successfully employed a comprehensive molecular dynamics framework to systematically evaluate the structural mimicry potential of three gut microbiota-derived peptides (KP1, KP2, and KP3) relative to a human self-peptide (ANX). Through integrated analysis of backbone RMSD, per-residue RMSF, SASA, radius of gyration, H-bonding networks, and MM-GBSA BE, we identified distinct mimicry phenotypes among the microbial peptides. Peptide KP1 emerged as a strong molecular mimic, which exhibits similar functional behaviour as the self-peptide across all structural and energetic descriptors. Its stable conformation, similar to ANX, sustained hydrogen bonding, and favourable binding thermodynamics collectively suggest it as a strong mimicry candidate for the HLA-B allele and possibly act to induce TCR cross-reactivity.

In contrast to this, KP2 demonstrated significant structural and energetic deviations from the reference peptide, which indicates minimal mimicry potential, whereas Peptide KP3 presented as an intermediate mimicry candidate by favourable energetics but conformational instability, suggesting it as a transient rather than sustained mimicry candidate. Our findings demonstrate the value of multi-parameter computational screening for prioritising mimicry candidates before expensive and time-intensive experimental validation. Comparing microbial peptides sequence similarity along with functional similarity via using multiple descriptors, as used in the present study, can serve as quantitative criteria for future computational screens of pathogen-derived epitopes.

## 5. FUTURE PERSPECTIVES

The computational framework presented here establishes a foundation for high-throughput screening of molecular mimicry candidates. Future investigations will extend this workflow to larger peptide libraries derived from diverse microbial sources, enabling genome-scale identification of potential autoimmune triggers. The integration of this multi-parameter MD analysis pipeline with high-throughput docking protocols and machine learning-based affinity prediction could rapidly filter thousands of peptide sequences to prioritise the most promising mimics for focused structural analysis. Such scalability is essential for comprehensive microbiome-autoimmunity studies where hundreds of candidate peptides must be evaluated. To directly assess T-cell receptor cross-reactivity, subsequent work should additionally incorporate explicit TCR modeling to create full peptide-MHC-TCR ternary complexes. Moreover, the future work will also include extended and replica simulations to improve sampling, as well as advanced analyses such as principal component analysis and conformational clustering to identify dominant motion patterns and stable structural states.

Furthermore, an exploratory machine learning exercise was tested in the present workflow outside the current project scope to generate a predictive model with a very limited data set, which can be regenerated with a longer data set and by adding more analytical properties. This predictive model can be used to predict a peptide’s initial mimicry potential without taking longer simulation steps. Finally, the computational predictions must be validated through rigorous experimental assays and cross-reactivity data from T-cell activation assays. By pursuing the suggested multidisciplinary research directions, molecular mimicry can be transformed from a mechanistic hypothesis into a quantitative, predictive framework for understanding and ultimately preventing microbiome-triggered autoimmune diseases.

## 6. Author Contributions

**Sanju Singh**: writing - original draft, review and editing, conceptualisation, investigation, visualisation, validation, methodology, software, formal analysis, resources, data curation.

## Acknowledgements

The author gratefully acknowledges the access to computational resources available during the research visit at Delft University of Technology (TU Delft), the Netherlands.

## 8. Reproducibility and Data Availability Statement

All scripts used for peptide generation, filtering, molecular dynamics simulations, and post-simulation analyses are publicly available in the project’s GitHub repository ^44^ which can support the reproducibility of the workflow and presented in the current study.

