## Supplemental Tables for "Molecular Dynamics Analysis of Self and Microbial Peptides Bound to HLA-B Protein: A Multi-Parameter Framework"

Sanju Singh*

**Supporting Information**

**Table S1.** MM-GBSA Energy Component Decomposition

| **Energy Component** | **ANX** | **KP1** | **KP2** | **KP3** |
| --- | --- | --- | --- | --- |
| ΔVDWAALS (kcal/mol) | -92.20 | -73.65 | -51.29 | −70.5 |
| ΔEEL (kcal/mol) | -568.35 | -422.65 | -195.55 | −415.4 |
| ΔEGB (kcal/mol) | 570.51 | 420.09 | 217.55 | 405.9 |
| ΔESURF (kcal/mol) | -15.45 | -13.09 | -8.32 | −8.9 |
| ΔGGAS (kcal/mol) | -660.55 | -496.30 | -246.83 | -485.9 |
| ΔGSOLV (kcal/mol) | 555.06 | 407.00 | 209.23 | 397.0 |
| **ΔTOTAL (kcal/mol)** | **-105.5 ± 7.2** | **-89.3 ± 20.3** | **-37.6 ± 10.7** | **-88.9 ± 18** |
| **ΔΔG vs anx (kcal/mol)** | **0.0** | **16.2** | **67.9** | **16.6** |

Standard deviations shown for ΔTOTAL are propagated errors calculated from component variances. ΔVDWAALS: van der Waals contribution; ΔEEL: electrostatic contribution; ΔEGB: polar solvation (Generalized Born); ΔESURF: non-polar solvation; ΔGGAS: gas-phase interaction energy; ΔGSOLV: solvation free energy. Calculations performed using gmx_MMPBSA with igb=5 and salt con=0.150 M.

**Table S2.** Key Per-Residue Energy Contributions from MM-GBSA Decomposition Analysis

| **Residue** | **ANX** | **KP1** | **KP2** | **KP3** |
| --- | --- | --- | --- | --- |
| **HLA Protein Residues:** |  |  |  |  |
| Glu64 (A-chain) | -5.87 | -3.04 | -3.22 | -3.5 |
| Arg63 (A-chain) | -1.54 | 0.06 | -3.02 | -0.5 |
| Glu77 (A-chain) | -0.80 | -0.63 | -0.11 | -0.7 |
| Lys147 (A-chain) | -5.07 | -8.73 | -0.62 | -7.5 |
| Trp148 (A-chain) | -4.64 | -2.84 | -0.57 | -3.2 |
| Tyr60 (A-chain) | -1.30 | -0.09 | -0.07 | -0.6 |
| Glu153 (A-chain) | -0.75 | -0.20 | -0.10 | -0.5 |
| **Peptide Residues:** |  |  |  |  |
| Position 1 (Ile/Gly/Gly/Ser) | +3.48 | +10.13 | +12.37 | +8.5 |
| **Position 2 (Arg/Arg/Arg/Arg)** | **-10.14** | **-10.54** | **-11.09** | **-10.8** |
| Position 3 (Ser/Ser/Ser/Ser) | -1.65 | -1.86 | -2.13 | -1.9 |
| Position 4 (Glu/Asp/Asp/Asp) | -0.73 | +1.03 | +1.98 | +1.2 |
| Position 5 (Phe/Phe/Phe/Arg) | -6.06 | -5.00 | -2.71 | -4.8 |
| Position 6 (Lys/Lys/Lys/Val) | +0.13 | -2.38 | -0.86 | -1.8 |
| Position 7 (Arg/Gly/Gly/Ala) | -4.63 | -0.76 | +0.09 | -1.5 |
| Position 8 (Lys/Asp/Asp/Lys) | -2.24 | -2.97 | +0.79 | -2.5 |
| **Position 9 (Tyr/Tyr/Tyr/Tyr)** | **-13.27** | **-10.42** | **-1.58** | **-11.5** |
| Position 10 (—/—/Phe/—) | — | — | -2.37 | — |

Values represent total binding energy contribution per residue (sum of van der Waals, electrostatic, and solvation components) in kcal/mol. Negative values indicate favorable (stabilizing) contributions; positive values indicate unfavorable (destabilizing) contributions. Only residues contributing |ΔG| > 0.5 kcal/mol are shown. Bold entries highlight critical anchor positions (Position 2: Arg; Position 9: Tyr).
